# Nematodes Vector Bacteriophages in Compost and Soil

**DOI:** 10.1101/2024.08.02.605737

**Authors:** Lisa van Sluijs, Cassidy Dietz, Floris van Noort, Johannes Helder, Mark W. Zwart, Kyle Mason-Jones

**Affiliations:** Wageningen University and Research, Laboratory of Nematology, Netherlands; Netherlands Institute of Ecology, Department of Terrestrial Ecology, Netherlands; University of New Hampshire, Department of Natural Resources and the Environment, New Hampshire, United States; Netherlands Institute of Ecology, Department of Microbial Ecology, Netherlands

**Keywords:** bacteriophages, nematode vectoring, hitchhiking, dispersal, phage-host interactions

## Abstract

Bacteriophages (phages) infect bacteria to reproduce and are often lethal to their host. To maintain the infection cycle, soil phages must travel from one suitable host to the next. However, traversing the soil matrix presents a dangerous journey for phages, which need to encounter the right hosts in this spatially complex habitat, but are not capable of active motion and can adsorb to soil particles. Here, we tested the hypothesis that bacterial-feeding nematodes (roundworms) present a reliable vehicle of soil transport for phages. First, we demonstrated that the bacterivorous nematode *Caenorhabditis elegans* vectored the laboratory model phage T7 as well as the soil phage Φ Ppu-W11 on agar. Sorption assays carried out with paralyzed and non-paralyzed nematodes showed that phage transport can occur via both external cuticular attachment and ingestion, and that the presence of host bacteria is not required for phage vectoring. Finally, we designed a microcosm to test phage transfer in compost and sandy soil using *C. elegans* and its sister species, *C. remanei*, respectively. This experiment confirmed that nematodes also enable phage movement through complex spatial habitats. This novel mechanism of phage vectoring extends our understanding of virus transmission in soil, revealing new multitrophic interactions that may influence soil functioning.

## Introduction

Bacteriophages (phages), viruses which infect bacteria, are the most abundant biological entities on earth, present in every habitat from the ocean to the mammalian gut (Suttle, 2007; Clokie et al., 2011; Williamson et al., 2017). Phages are also highly prevalent in soil and are likely a major driver of bacterial death in terrestrial ecosystems (Thurber et al., 2017), thereby enhancing nutrient and carbon cycling (Ashelford et al., 2003; Williamson et al., 2017; Emerson et al., 2018; Kuzyakov and Mason-Jones, 2018). However, to maintain the infection cycle, phages need to disperse from one host to the next, despite having no independent metabolism or motility. This is a particularly daunting challenge in complex soil habitats.

Soils represent a number of difficulties for phage dispersal: (i) bacteria are sparsely and patchily distributed in soil (Baveye et al., 2018); (ii) phages have a narrow host range, while soil bacterial populations are highly diverse; (iii) soil minerals, organic matter, and liquid interfaces can sorb or damage phage particles (Jin and Flury, 2002); and (iv) the aqueous phase through which phages could diffuse is highly discontinuous (Erktan et al., 2020). Nonetheless, extracellular phages must travel through this soil labyrinth to reproduce. Random diffusion is slow and undirected, while mass flow during wetting events is occasional and unreliable. In view of these challenges, the abundance of extracellular phage particles in soil suggests that other mechanisms may help them to encounter their host bacteria (Nimmo, 2004; Young et al., 2008; Dennehy, 2014).

Pioneering evidence shows that microbes, including phages, may travel by attachment to motile organisms, referred to as hitchhiking. Non-host bacteria travelling along fungal hyphae can co-transport phages even across air-filled voids (You et al., 2022b, 2022a). Nematodes, commonly referred to as roundworms, have been found to transport bacteria (Thutupalli et al., 2017) and phages in pure culture (Dennehy et al., 2006). Additionally, some plant-feeding nematodes are well-known vectors of plant viruses (MacFarlane and Robinson, 2001). In soils, bacterivorous nematodes can be highly abundant (densities up to 10 million per m^2^) and actively search for bacterial prey with the benefit of sensory navigation (Moens et al., 1999; Choi et al., 2016; Liu et al., 2017; van den Hoogen et al., 2019; Chai et al., 2024). Together, this suggests a potential for nematodes to act as phage vectors, but this has not been previously investigated in spatially structured habitats such as soil and compost, nor with native soil phages.

Here we examined whether nematodes could vector phages between patches of host bacteria, and how this might occur. First, we examined whether the nematode *Caenorhabditis elegans* was essential for movement of the laboratory model phage T7 and the soil phage Φ Ppu-W11 on agar. Next, we performed sorption assays with paralyzed and non-paralyzed nematodes to test probable mechanisms of this transport: transport of phage particles versus transport of phage-infected hosts; and adhesion to the nematode cuticle (outer surface) versus ingestion and excretion. Finally, we conducted a microcosm experiment to test phage transfer in compost (with *C. elegans*) and a sandy soil (with a *C. remanei* isolate from soil), thereby for the first time testing phage transfer by nematode hitchhiking in complex spatial habitats.

## Methods

### Experimental organisms and culture conditions

The experiments were conducted with *Escherichia coli* REL606 and OP50 (Brenner, 1974; Jeong et al., 2009) the *E. coli* phage T7, *Pseudomonas putida* and the *Pseudomonas* phage Φ Ppu-W11. The latter two were both obtained from the Leibniz Institute German Collection of Microorganisms (accession DSM291 and DSM100071, respectively). *P. putida* is a widespread soil bacterium, and Φ Ppu-W11 was isolated from soil in Germany in 2014 (DSMZ, 2014). *P. putida* cultures were prepared from a single active colony suspended in 5 mL tryptic soy broth (TSB) medium with 5 mM MgSO_4_ and shaken (160 rpm, 27℃) overnight prior to use in experiments. *E. coli* cultures were prepared from a single active colony added to 5 mL Luria broth (LB) medium with 5 mM MgSO_4_ and shaken (160 rpm at 37℃) overnight prior to use. Phage lysates were prepared by adding 200 µL of host overnight culture and 50 µL of phage stock to 5 mL of fresh LB (*E. coli*) or TSB (*P. putida*) liquid medium, then shaking (160 rpm at 37℃ or 27℃ respectively) for approximately 6 hours until lysis was evident in clearing of the culture medium. Lysed culture was centrifuged (10 min at 4000 ×g) and syringe filtered to 0.22 µm. Lysate was stored at 4°C.

WN2002 (WBStrain00040481) is a wild isolate of the bacterivorous nematode *Caenorhabditis elegans*, collected from a garden compost heap in Wageningen, The Netherlands (Cook et al., 2017; Crombie et al., 2024). N2 is the *C. elegans* reference strain, originally isolated from mushroom compost in Bristol, United Kingdom. *C. remanei* strain WN210S was collected from a sandy garden soil in Wageningen (gifted by Guixin Li). Nematodes were cultured at 20 ℃ (*C. elegans*) or 16 °C (*C. remanei*) on 6 cm petri dishes with nematode growth medium (NGM) (Brenner, 1974) pre-seeded with 50 µL *P. putida* or *E. coli* OP50 overnight culture. Nematode strains for experiments were maintained via chunking (agar transfer) every few days and were harvested for experiments by suspension into M9 buffer between 3 and 5 days after chunking (Brenner, 1974; Meneely et al., 2019). Resulting nematode suspensions contained mixed life stages (eggs, L1, L2, L3, L4, adult) and some bacteria from the culture plate. When required, nematodes were sterilized from bacteria by bleaching to obtain sterile eggs (Meneely et al., 2019).

### Phage enumeration

Phage abundance in lysates and extracts from agar or soil was quantified via soft-top plaque assays on TSB (*P. putida*) or LB agar (*E. coli*) (Clokie and Kropinski, 2009). Briefly, serial dilutions of phage lysate or extract (100 µL) and overnight culture of the respective host (100 µL) was mixed with 7 mL of soft agar (7 g L^-1^, 55 °C), and immediately poured over a petri dish half-filled with agar, followed by overnight incubation at the respective growth temperature. Host bacteria grow to form a solid lawn, except where phage lysis prevents this, leaving translucent gaps (plaques). Phage abundance was calculated as plaque forming units (pfu’s) from plaque counts.

### Nematode hitchhiking on agar

Phage transfer by *C. elegans* nematodes across an agar surface was tested in a setup similar to Dennehy *et al*. 2006 (Figure 1). For the *P. putida* experiment, resource heterogeneity was introduced to 9 cm petri dishes containing nematode growth medium agar by removing two 2 cm-diameter plugs from the agar, 2 cm apart, using a flame-sterilized corer and refilling the holes with 0.1X and 1X TSB agar, respectively. The 0.1X TSB provided a low resource patch and 1X TSB a high resource patch. 20 µL *P. putida* overnight culture was inoculated onto each TSB patch. For the *E. coli* experiment, 20 µL overnight culture was directly spread onto a patch of the NGM agar. Nematode suspensions were obtained from nematode culture plates (therefore also including some *E. coli* or *P. putida* from the culture plate, corresponding to the host of the phage). A 20 µL *C. elegans* nematode suspension containing 20-200 nematodes, with or without 10 µL phage lysate (titer T7: 2.5·10^4^, Φ Ppu-W11: 1·10^7^ pfu/mL) was spread on an agar plate at a point 2 cm from the one (*E. coli*) or two (*P. putida*) bacterial patches. All plates were incubated in the dark at 20°C for 40 hours.

**Figure 1.**
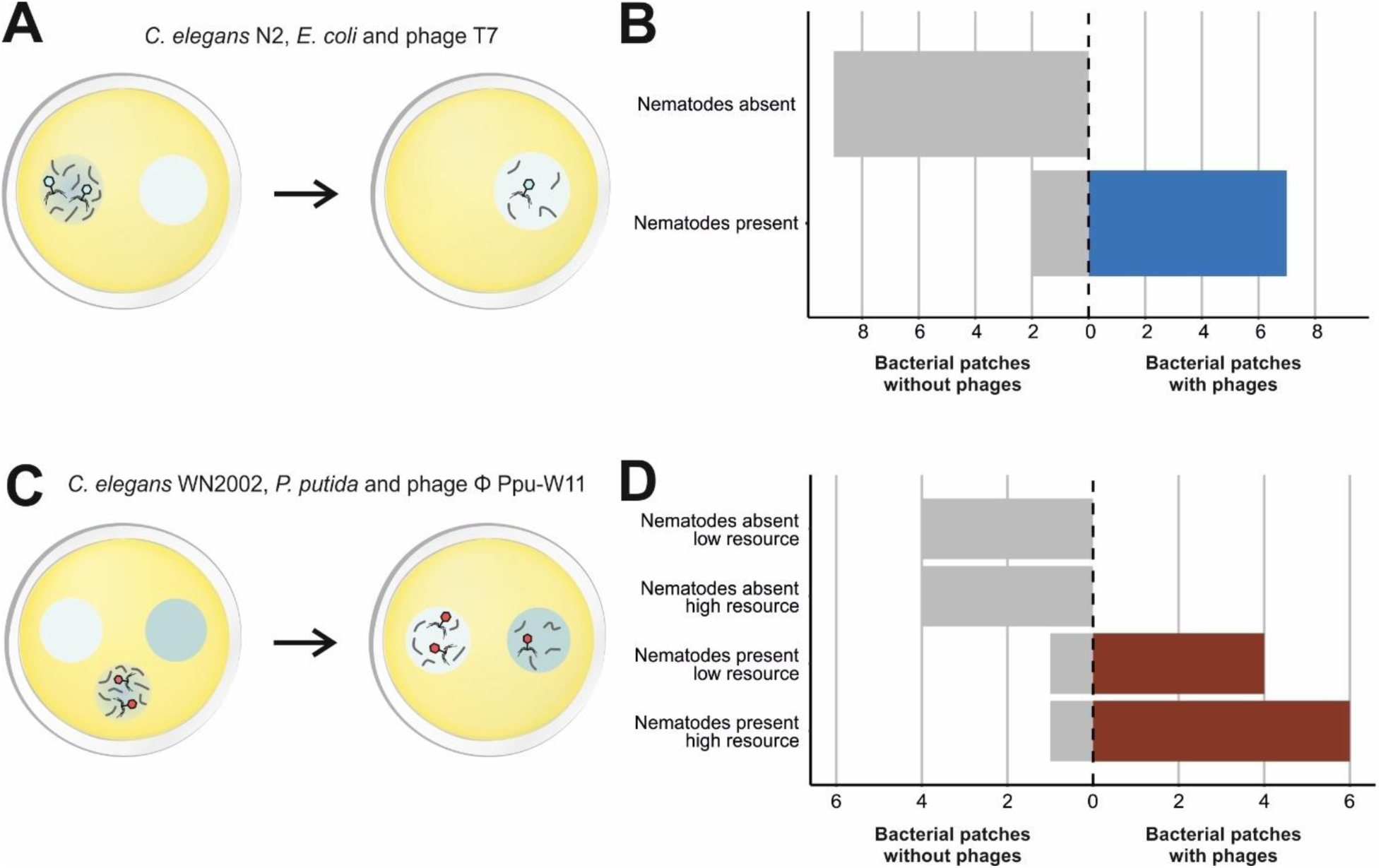
Nematode hitchhiking on agar. – A) Nematode *C. elegans* strain N2 and *E. coli* phage T7 were placed on agar 2 cm away from a patch of *E. coli* host bacteria. Control plates were identical, except lacked nematodes. B) Presence of phage T7 in bacterial patches 40 hours after inoculation was dependent on the presence or absence of *C. elegans* nematodes in the patch (n = 9). The number of bacterial patches (horizontal axis) represents the total number of experimental observations where presence (phage transfer) or absence (no phage transfer) of phages was observed in a bacterial patch. C) *C. elegans* strain WN2002 and *Pseudomonas* phage Φ Ppu-W11 were added 2 cm away from two patches of *P. putida* host bacteria, with contrasting resource availability. D) Presence of Φ Ppu-W11 phage in the high and low resource patches when *C. elegans* nematodes were absent or present in the patch (nematodes absent: n = 8; nematodes present: n = 12, with n as the number of bacterial patches). The number of bacterial patches (horizontal axis) represents the total number of experimental observations where presence (phage transfer) or absence (no phage transfer) was observed in a bacterial patch.

After incubation, nematode presence in each bacterial patch was confirmed under the microscope (Leica M205C at magnification 17.5-25×). Thereafter, ∼1 cm of the bacterial patches were removed with a flame-sterilized spatula into 8 mL phage extraction buffer (BSA-supplemented PBS; (Göller et al., 2020)) and shaken (70 rpm) for 2 hours at 4 ℃ before centrifugation (5 minutes at 9400 ×g) and syringe filtering to 0.22 µm. The phage titres of these extracts were enumerated by soft-top assay. Phage titres represent both transferred phages as well as phages produced through new infections in the bacterial patch or during extraction, and therefore provide a sensitive binary measure of phage transfer (presence/absence) but not a quantification of phages transferred. Some results were excluded from the data as soft-top assays failed (Supplementary Table S1).

### Nematode vectoring in the absence of bacterial hosts

To examine whether nematodes can vector phage particles in the absence of their bacterial host, *P. putida* overnight culture (20 µL) was spread using a flame-sterilized loop onto a 1.5 cm-diameter patch on one half of a 9 cm TSB agar plate and incubated at 20 °C overnight. The following day, cell-free *P*. *putida* phage lysate (20 µL, titer: 2·10^9^ pfu/mL) was inoculated in a 0.75 cm wide strip across the middle of each plate (Figure 2). Nematode suspensions were prepared from nematodes grown on *E. coli* OP50 (non-host for Φ Ppu-W11 phage) and pipetted onto a spot diametrically opposite the incubated patch of *P. putida*, separating *C. elegans* nematodes and bacteria by the strip of phage. All microcosms were incubated in the dark at 20℃ for 2 hours.

**Figure 2.**
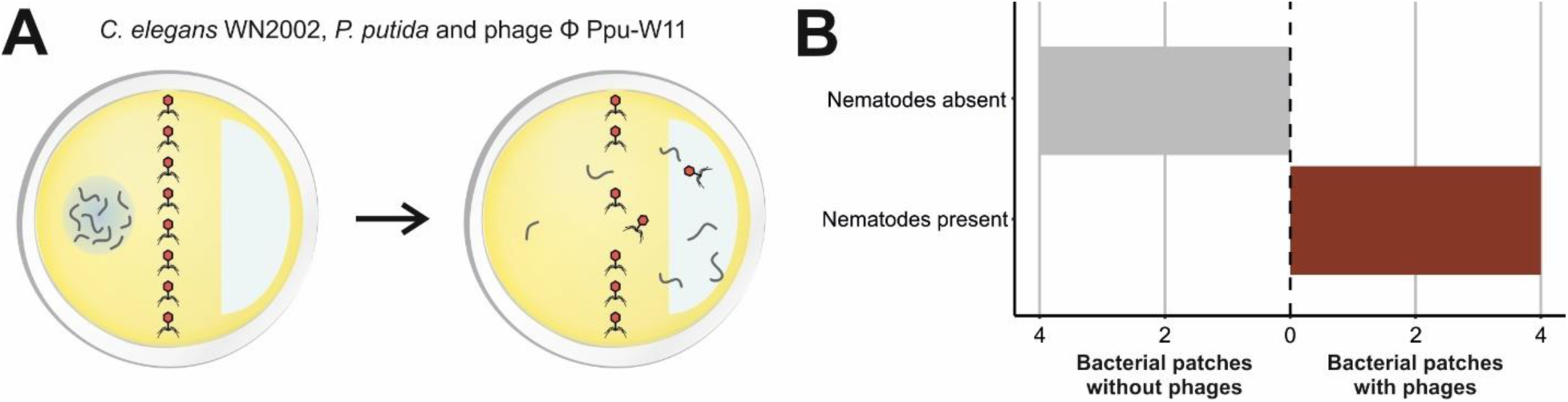
Crossing the line: ‘naked’ phage transfer by nematodes. – A) *C. elegans* WN2002 (grown on non-host *E. coli*) was added on one side of an agar plate with host *P. putida* inoculated on the other side of the plate. These were separated by a line of *Pseudomonas* phage Φ Ppu-W11 cell-free suspension. Phages were extracted from the *P. putida* patch after 2 hours of incubation. B) Presence of Φ Ppu-W11 phage in the bacterial patches when *C. elegans* nematodes were absent or present (n = 4). The number of bacterial patches (horizontal axis) represents the total number of experimental observations where presence (phage transfer) or absence (no phage transfer) was observed in a bacterial patch.

After incubation, nematodes on *P. putida* patches were counted under the microscope. The host patches were excised from the agar using a flame-sterilized spatula and extracted with 5 mL of TSB liquid medium in a 50 mL centrifuge tube (27 °C with shaking at 170 rpm) in order to amplify the likely small amount of vectored phage to detectable levels, and then used in a soft-top assay (Göller et al., 2020).

### Mechanisms of nematode vectoring

*C. elegans* WN2002 were grown on *E. coli* OP50. Nematodes (∼250 individuals of mixed life stages) were suspended in M9 buffer with or without 1 mM levamisole (a nematode paralysing agent) along with 10 µL *P. putida* phage Φ Ppu-W11 (titer: 1·10^8^ pfu/mL) in presence of either host *P. putida* (35 µL overnight culture) or non-host bacteria *E. coli* REL606 (35 µL overnight culture) (Figure 3). Blockage of nematode feeding by 1 mM levamisole was confirmed after visualisation of feeding on fluorescent beads as described in previous studies (Bakowski et al., 2014; van Sluijs et al., 2021). This phage exposure to Φ Ppu-W11 was performed in a mixed, total volume of 500 µL for half an hour at room temperature (∼20 °C), including an additional mixing step after 15 minutes. After exposure, all samples were washed thrice with 1 mL of M9 buffer including 1 mM levamisole to remove bacteria and unabsorbed phages. Then the supernatant containing the nematodes was lysed using a Ø3mm steel bead in a QiaGen TissueLyser (5 min at 30 Hz). Phages were extracted with 100 µL BSA-amended PBS extraction buffer and quantified by soft-top plaque assay (Göller et al., 2020). Nematode-free control samples (same compositions except for the nematodes) were processed identically. In some cases, plates contained too many plaques to quantify transfer (confluent or semiconfluent lysis). In these cases, additional experimental replicates were performed using a diluted stock of phage Φ Ppu-W11 (titer: 1·10^6^ pfu/mL) (Supplementary Table S3). Phage titres represent transferred phages when non-host *E. coli* REL606 was used. With *P. putida*, phage infections may have also contributed to phage titres, and the short period of host exposure (30 minutes) and direct quantification by soft-top plaque assay were used to limit this bias.

**Figure 3.**
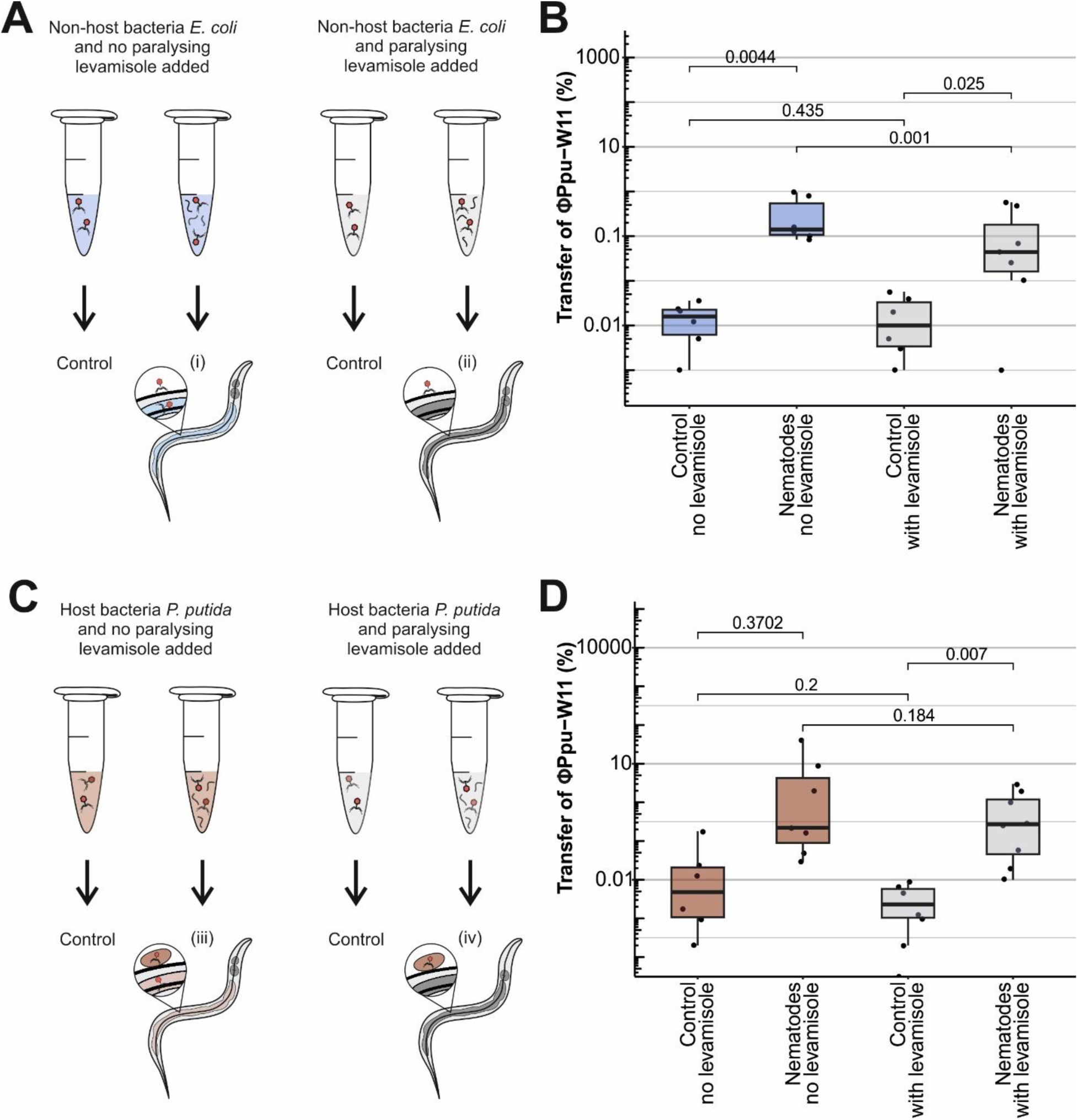
Mechanisms of nematode hitchhiking by phages. – Nematode *C. elegans* WN2002 was incubated for 30 minutes with bacterial prey and *Pseudomonas* phage Φ Ppu-W11 in either presence or absence of the nematode paralysing agent levamisole. Without levamisole, bacteria or phages could be attached to the nematode external cuticle or have been ingested. With levamisole nematodes cannot feed, so only externally attached phages or bacteria are present after washing. Control samples lacked nematodes A) Non-host *E. coli* as prey. Phage particles associated with active nematodes after washing could be attached to their external cuticle or have been ingested, but not as infected bacteria. With levamisole only externally attached phage particles are expected. B) Percentage (out of the total added to the solution) of phages associated with nematodes in the presence of non-host *E. coli* (n = 8). C) Phage host *P. putida* as prey, enabling attachment or ingestion as infected bacteria; D) Percentage (out of the total added to the solution) of phages associated with nematodes in the presence of phage host *P. putida* (n = 6).

### Nematode vectoring in compost and sandy soil

Phage transfer was assessed in compost using *C. elegans*, *P. putida* and *Pseudomonas* phage Φ Ppu-W11. Organic compost (Pokon Naturado B.B., The Netherlands) was sterilized by two rounds of autoclaving (121℃, 30 mins, 24 h apart). Compost microcosms were constructed in glass troughs with rectangular cross-sections 10 mm wide by 12 mm high (Figure 4). Aliquots of *P. putida* overnight culture (5 mL) were centrifuged (9400 ×g, 10 min) and the pellets were resuspended in either Tris phage buffer (low resource) or TSB liquid medium (high resource). These *P. putida* suspensions (100 µL/g) were added to separate aliquots of the sterile compost, thoroughly mixed, and packed onto the ends in 1 cm-long segments. *C. elegans* (WN2002) nematodes were obtained from 3-day-old cultures by suspending nematodes in M9 solution. Each microcosm was inoculated in the middle of the trough with 30 µL of either nematode suspension (approximately 1600 *C. elegans* nematodes) or M9 buffer (non-nematode control) and with *P. putida* Φ Ppu-W11 (1·10^6^ pfu), leaving 2 cm of sterile compost separating the inoculation segment from the bacterial segments at the ends. All segments of compost were separately prepared and to reach a uniform moisture content of 100% dry wt. after all inoculations.

**Figure 4.**
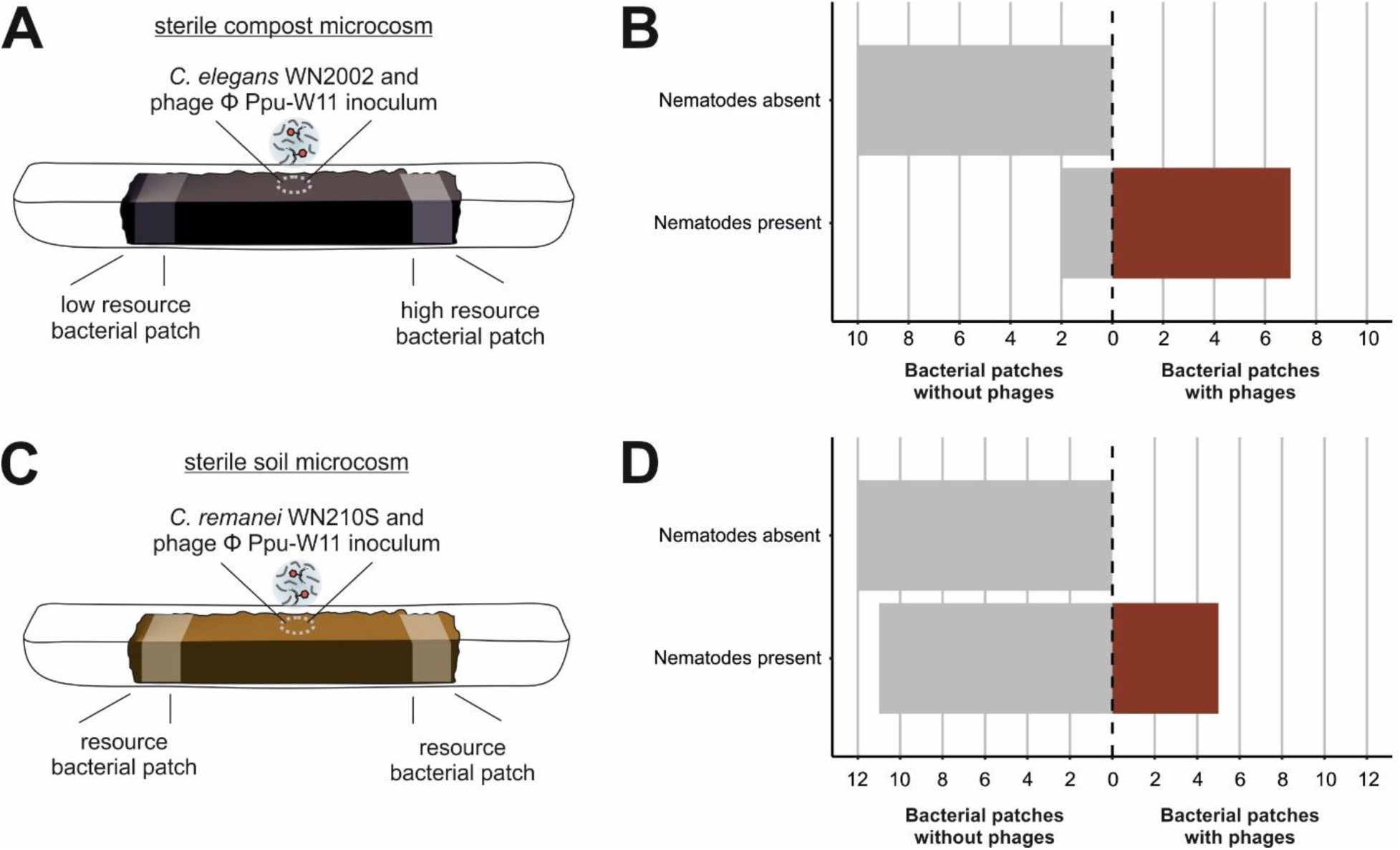
Phage transfer by nematodes in spatially structured habitats. – A) Sterile organic compost was flanked by two sections with host *P. putida* inoculated in either high or low resource medium. *Pseudomonas* phage Φ Ppu-W11 and *C. elegans* WN2002 were then introduced to the middle of the compost and phage and nematode presence in the outer bacterial patches was assessed after 3 days incubation. Control conditions lacked nematodes. B) Observed phage vectoring in A in relation to the presence or absence of nematodes in outer bacterial patches (nematodes present: n = 9; nematodes absent: n = 10, with n as the number of bacterial patches). The number of bacterial patches (horizontal axis) represents the total number of experimental observations where presence (phage transfer) or absence (no phage transfer) of phages was observed in the bacterial patch. C) The same setup was constructed with sterile sandy soil but with a soil isolate of the nematode *C. remanei* WN210S and only high resource patches. D) Observed phage vectoring in C in relation to the presence or absence of nematodes in outer bacterial patches (nematodes present: n = 16; nematodes absent: n = 12, with n as the number of bacterial patches). The number of bacterial patches (horizontal axis) represents the total number of experimental observations where presence (phage transfer) or absence (no phage transfer) of phages was observed in the bacterial patch.

A sandy soil (85% sand, 9% silt and 6% clay) was sampled at Wageningen University and Research’s experimental farm to a sampling depth of 20 cm, forming a composite sample of 10 cores on a field transect. The soil contained 1.1% organic C and had a pH of 6.2. Sandy soil microcosms were constructed in the same way as for compost, but with a *C. remanei* nematode (30 µL suspensions containing approximately 400 *C. remanei* nematodes) strain isolated from a comparable sandy soil; *P. putida* was only added in TSB liquid medium (high resource); and the final moisture content was a uniform 19% dry wt.

All microcosms were wrapped in autoclaved aluminium foil and incubated at 20°C without light for 3 days. Bacterial patches were then harvested by pulling them free of the main soil body and gently mixing within the glass trough. Nematodes were extracted by transferring approximately 100 mg of compost or sandy soil from a bacterial patch onto a plate inoculated with *P. putida* so that nematodes would leave the compost to forage. After overnight incubation (2 0℃), nematodes that had migrated out onto the agar surface were counted under the microscope. 0.5 g compost samples from each resource patch were extracted in 10 mL BSA-amended PBS and shaken (50 rpm) for 1 hour at 4℃, centrifuged (10 minutes at 9400 ×g), and syringe filtered to 0.22 µm, prior to soft-top plaque assays.

### Data analysis

Analyses were performed in R (version 4.3.2) using package “tidyverse” (Wickham et al., 2019; R Core Team, 2020). All experiments, except for the mechanistic sorption experiment, used a host population as a nematode attractant and infection in this population as an indicator of phage arrival. Since a single phage infection in a dense population can rapidly generate large numbers of new phages, this approach provides a highly sensitive presence/absence measure but not a quantitative measure of the number of phages transferred. Data were therefore analysed based on absence or presence of phage using Firth’s logistic regression with binomial distribution (R package “logistf”, (Heinze et al., 2023)). For the compost microcosm experiment, low and high resource data was combined, as no significant effect of resource level was observed. Data from the mechanistic sorption experiment were analysed with a linear mixed effects model (LMM) using restricted maximum likelihood (REML) (R package “lme4”, (Bates et al., 2020)) with treatment (with/without nematodes and levamisole) as fixed effect and a random batch effect according to the following model:

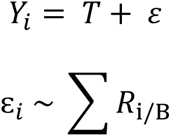

where Y is the recovery of the phage (%) that was explained over treatment (T), and the error term ε based on the autocorrelation matrix of technical replicate R_i_ (1, 2,…, 11) given biological replicate B (1, 2,…, 5). The *Anova* function of the “car” package was used to compute the analysis of variance (Fox and Weisberg, 2019). Results from statistical analyses are included in Supplementary Table S5.

### Data availability

*C. elegans* strains used in this study are available from CaeNDR (caendr.org) and the Caenorhabditis Genetics Center (Cook et al., 2017; Crombie et al., 2024). Custom written scripts and data files can be found on https://git.wur.nl/published_papers/Phage_transfer_by_nematodes. Processed data is included in the Supplementary Tables of this manuscript.

## Results

### Nematodes vector model and soil phages on agar

When foraging on agar, phage-exposed *C. elegans* nematodes demonstrated consistent vectoring of phages to uninfected bacterial patches (Figure 1). This was the case for both the laboratory model phage T7 and the soil isolate of *Pseudomonas* phage Φ Ppu-W11. Almost without exception, the presence of nematodes in a bacterial patch was accompanied by phages, and no phage transfer occurred without nematodes, confirming that nematodes were essential for phage transfer (Firth’s logistic regression: p_T7_ = 6.2 · 10^-4^, p_Φ Ppu-W11_ = 1.9 · 10^-4^) (Figure 1B, D, Supplementary Table S1, S5). With *P. putida*, we additionally tested whether a higher bacterial density would attract more nematodes, but nematodes showed no preference between high- and low-density patches (Supplementary Table S1) (Firth’s logistic regression: p = 0.78). These results indicate that both the *E. coli* phage T7 and the soil phage Φ Ppu-W11 are vectored by nematodes.

### Phage vectoring not reliant on host presence

Next, we investigated the mechanisms by which nematodes vector phages, by asking whether the ingestion of bacteria was a requirement for phage transfer. We tested whether nematodes that crawled through a strip of cell-free Φ Ppu-W11 would subsequently vector phage into a patch of *P. putida* (Figure 2A). Nematodes arrived in the *P. putida* patches within 2 hours of inoculation (7-53 nematodes per patch). Phages were transmitted in every plate where nematodes were present (4/4 replicates) (Figure 3B, Supplementary Table S2), verifying that phage was vectored from the host-free environment to the host patch (Firth’s logistic regression: p = 6.7 · 10^-3^). Moreover, phage transmission was not detected when nematodes were absent. Concluding, we therefore find indicative evidence that nematodes can vector phages in the absence of the host bacterium and without active feeding.

### Nematodes vector phages via cuticle adhesion and ingestion

We hypothesized four mechanisms by which nematodes may vector phages: (i) ingestion and excretion of phage particles (Figure 3A); (ii) external attachment of phage particles (Figure 3A); and (iii) ingestion and excretion of infected host (Figure 3C) or (iv) external attachment of infected host (Figure 3C). Paralysis of the nematodes caused a 2.5-fold reduction in nematode-associated phages in the presence of non-host *E. coli* (Figure 3B), reflecting mechanism (i) (LMM, p = 0.001). Supporting mechanism (ii), presence of nematodes paralyzed by levamisole increased phage recovery compared to the nematode-free control (LMM, p = 0.025) (Figure 3B, D). Phage recovery percentages of paralyzed nematodes were higher in host than non-host presence (LMM, p = 0.008) (Supplementary Table S5), thus host presence (iv) may further increase vectoring (Figure 3D), but we cannot rule out the possibility of increased phage counts due to replication in the host, despite the short (30 min) exposure. Contrary to mechanism (iii), nematodes exposed to *Pseudomonas* phage and its *P. putida* host while active or paralyzed were associated with similar numbers of phages (Figure 3D) (LMM, p = 0.184). Together, these results indicate that internal and external hitchhiking of phage particles can both contribute to vectoring. Moreover, although we cannot conclude that hosts enhance phage transfer (because of potential phage replication in the host bacteria), it is clear that hosts are not necessary for this transfer.

### Nematodes vector phages across complex compost and soil environments

We tested whether nematodes vector phages in complex environments with a novel set-up constructed from sterile compost or soil (Figure 4A, C, respectively). In the absence of nematodes, no phages were found in bacterial patches 2 cm from the phage inoculation site (Figure 4), while phage transfer did occur when nematodes were co-inoculated with the phages (Firth’s logistic regression: p = 3.9 · 10^-4^) (Figure 4B, Supplementary Table S4). *C. elegans* vectored phage across compost with high consistency, with phages detected in 78% of bacterial patches when nematodes were present (Figure 4B). *C. elegans*, a nematode associated with rotting substrates rather than soil (Barrière and Félix, 2005; Felix and Braendle, 2010; Frézal and Félix, 2015; Schulenburg and Félix, 2017), did not move through sandy soil in a pilot experiment. An isolate of the sister species *C. remanei* was found to move and vector phages through the sandy soil (31% of cases with nematode presence, Firth’s logistic regression: p = 3.6 · 10^-2^) (Figure 4D, Supplementary Table S4). These results conclusively demonstrate that nematodes can vector phages through compost and soil habitats, across distances of at least centimetre scale.

## Discussion

Phages are abundant in soil (Williamson et al., 2017), despite this habitat presenting formidable barriers to their transmission. Here we have demonstrated that nematodes are effective vectors for phages, offering an alternative to independent extracellular transmission of phage particles. We confirmed that nematode hitchhiking by phages on agar, as previously reported (Dennehy et al., 2006), is also relevant for a soil phage, and was insensitive to the resources available for host growth. Finally, we for the first time demonstrated phage vectoring in complex compost and soil environments. Considering that nematodes are omnipresent in terrestrial habitats (van den Hoogen et al., 2019) our results reveal a potentially important driver of phage infection and bacterial mortality.

### Implications of vectoring mechanism

We assessed nematode hitchhiking by phages via external (cuticular) and intestinal mechanisms. Based on experiments using *E. coli* phage Φ6, it was previously suggested that phages are only transferred in the form of infected bacteria rather than extracellular phage particles (Dennehy et al., 2006). Contrary to this expectation, vectoring of cell-free phage Φ Ppu-W11 was demonstrated across agar. Although we did not observe nematodes crawling back after reaching the bacterial patch, we could not fully exclude nematodes crawled back. Yet, another independent experiment also showed that the soil phage (Φ Ppu-W11) associated with nematodes in the presence of non-host bacteria (*E. coli*), and together these experiments imply host bacteria are not needed for phage transfer. The contribution of external versus internal transfer could only be conclusively determined in the presence of non-host bacteria, whereas potential replication of Φ Ppu-W11 in the presence of hosts hinders unambiguous interpretation. Therefore, we conclude that both internal and external transfers of phage particles contribute to vectoring by nematodes, but the relative quantification remains to be determined in the presence of the host bacterium.

Phage vectoring by cuticular attachment implies that phages remain in contact with soil, where sorptive minerals may reduce phage transfer. Less phage vectoring was observed in sandy soil than in compost, despite nematode dispersal in both substrates. The reduced rate of phage transfer in sandy soil is therefore consistent with a substantial role of external binding as a vectoring mechanism. Phage transport after ingestion or transport within infected hosts may provide more protected transport conditions and be a relatively more important driver in sorptive soil environments. However, these comparisons between soil and compost were confounded with the use of a different *Caenorhabditis* nematode species, which may also have variable transfer rates. Although *Caenorhabditis spp.* are not typical soil nematodes (Barrière and Félix, 2005; Felix and Braendle, 2010; Frézal and Félix, 2015; Schulenburg and Félix, 2017), the *C. remanei* strain in this study was isolated from sandy soil and dispersed through the soil matrix, whereas this was not observed for *C. elegans*. Generalizing across habitats and taxa will unavoidably involve such confounded comparisons, and future studies may focus on additional (soil) nematode species and nematode-bacteria-phage interactions in their natural habit to resolve aforementioned questions.

### Ecological interactions in phage transfer by nematodes

Motile organisms, including moving patches of bacteria, growing fungi and nematodes, have all been proposed as mechanisms of phage transfer (Dennehy et al., 2006; Ping et al., 2020; You et al., 2022b, 2022a), but these previous studies have not demonstrated their operation in spatially structured soil or compost environments. Nevertheless, this growing body of evidence supports recent interest in vectoring interactions as drivers of soil function (Muok and Briegel, 2021). Nematodes have long been known to spread plant viruses and bacteria (Brown et al., 1995; MacFarlane, 2003; Thutupalli et al., 2017) and phages can now be added to the list of nematode-borne microorganisms. Moreover, nematodes themselves can be vectored by macroscopic organisms such as isopods and slugs (Félix and Duveau, 2012; Petersen et al., 2015; Schulenburg and Félix, 2017; Archer et al., 2020), potentially expanding the dispersal range of a hitchhiking phage. These layered interactions highlight the importance of movement as a driver of viral ecology in soil.

Distinct species of nematodes have different surface properties and can carry specific cuticular microbiomes (Davies and Curtis, 2011; Elhady et al., 2017). This may result in variable binding between phage and nematode taxa. We speculate that nematodes may also carry distinct viromes, with the ability to directly attach to suitable nematodes may be adaptive for virulent phages, allowing them to escape collapsing bacterial populations. In contrast, the vectoring of phages that compete for their bacterial prey is presumably detrimental to fitness but might be unavoidable. Nematodes do, however, have various adaptations to avoid adverse organisms, for example by detection of chemical and nucleic acid-based cues (Kaletsky et al., 2020; Ferkey et al., 2021). Whether they possess the ability to evade phages is an intriguing question. On the other hand, phages might benefit nematodes (Church et al., 2000; Höss et al., 2001; Silva-Sánchez et al., 2019). Tens of bacterial species were identified in the natural habitat of *C. elegans* that hinder growth or even actively kill nematodes (Samuel et al., 2016). Nematodes may profit from phages that infect such detrimental bacterial species.

The ecological implications of phage vectoring will depend on selectivity in feeding behaviour of the nematode. Phages typically have constrained host ranges, and vectoring thus relies on nematode movement between patches of suitable host bacteria. Some studies provide evidence of species-selective feeding by nematodes (Berg et al., 2016; Liu et al., 2017; Dirksen et al., 2020; Zhang et al., 2021), but nematode feeding preferences at the level of bacterial species are yet to be determined in soil (Schuelke et al., 2018; Guden et al., 2021; Vafeiadou et al., 2022). Earlier studies indicated that foraging behaviour also depends on bacterial cell densities (Madirolas et al., 2023). We did apply different bacterial densities, but did not observe evidence for differences in phage transfer resulting from foraging behaviour.

## Conclusions

We demonstrated that nematodes vectored soil phages across agar, soil and compost habitats over centimeter scale distances. This vectoring occurs also in the absence of host bacteria, indicating the transfer of extracellular phage particles. Vectoring mechanisms include both adhesion to the external cuticular surface of the nematode and ingestion followed by excretion. Overall, our study provides novel insight into how phages navigate complex soil habitats to infect new hosts by hitchhiking with nematodes. This invites further study of phage-nematode-bacteria interactions in terrestrial ecosystems and potential applications such as steering soil communities by addition of nematode-phage pairs.

## Acknowledgments

LvS was supported by the C.T. De Wit Production Ecology & Resource Conservation (PE&RC) collaboration grant. K.M.J. acknowledges the Dutch Research Council (NWO) for funding of the Veni project VI.Veni.202.086. *Pseudomonas putida* and *Pseudomonas* phage Φ Ppu-W11 were obtained from the Leibniz Institute German Collection of Microorganisms and Cell Cultures (DSMZ). Some strains were provided by the CGC, which is funded by NIH Office of Research Infrastructure Programs (P40 OD010440). We are grateful to Stefan Geisen and Guixin Li for sharing the *C. remanei* strain WN210S, and Slavica Ivanovic for laboratory support.

## CRediT authorship contribution statement

**Lisa van Sluijs**: Writing – Original Draft, Funding acquisition, Conceptualization, Formal analysis, Software, Investigation, Data Curation, Visualization, **Cassidy Dietz**: Writing – Original Draft, Investigation, **Floris van Noort**: Investigation, **Johannes Helder**: Conceptualization, Funding acquisition, **Mark Zwart**: Conceptualization, Funding acquisition, **Kyle Mason-Jones**: Conceptualization, Methodology, Supervision, Writing – Review & Editing, Funding acquisition.

